# A genome-wide approach uncovers the suite of genes important for swarming motility in the biocontrol bacteria *Pseudomonas protegens* Pf-5

**DOI:** 10.1101/2025.07.07.663420

**Authors:** BK Fabian, C Foster, A Asher, LDH Elbourne, KA Hassan, SG Tetu, IT Paulsen

## Abstract

Motility is a crucial bacterial trait, important for competitive success in many contexts. Plant growth promoting rhizobacteria, such as *Pseudomonas protegens* Pf-5, can protect agricultural crops against disease, relying on motility to access and colonize the plant rhizosphere. Swarming motility is the multicellular movement of bacteria powered by rotating flagella across a semi-solid surface which allows bacteria to move rapidly, search for and acquire food, avoid toxic substances, and colonize new niches. Here we performed genome-wide investigations, using transposon-directed insertion site sequencing (TraDIS) to identify 136 genes involved in *P. protegens* Pf-5 swarming motility. This method identified flagella biosynthesis, assembly, structure and regulatory genes, as expected. We also identified an additional suite of genes important for swarming, including chemotaxis, c-di-GMP turnover, cell division, peptidoglycan turnover, lipopolysaccharide biosynthesis, metabolism of thiamine (vitamin B_1_) and transport of pyoverdine. Overall, this study increases our understanding of pseudomonad motility and expands the breadth of lifestyles and habitats examined in relation to swarming motility. Understanding the genetic basis of swarming motility in biocontrol bacteria is important for successful application of these important plant growth promoting microbes in agricultural settings.

## Introduction

Plant growth promoting rhizobacteria (PGPR) can provide plants with benefits similar to fertilizer and pesticide application without many of the detrimental effects on the environment (1). One benefit biocontrol rhizobacteria can provide is protection from pathogens, but to do this they need to colonize the surface of the roots and the thin layer of soil around the roots, known as the rhizosphere (2, 3). Movement in the rhizosphere is essential for these bacteria to be able to colonize and provide protection (4, 5).

One type of motility important for rhizospheric bacteria is swarming. This is the multicellular movement of bacteria powered by rotating flagella across a semi-solid surface (6–9). Cells in swarming populations may differentiate to become longer and hyperflagellated (10–12). For example, *Pseudomonas aeruginosa* PA14 and PAO1 cells become elongated with multiple polar flagella when swarming (13, 14). The coordinated movement in swarming requires cell-to-cell signaling and the use of surfactants to reduce surface tension and drag (15). Pseudomonads employ a variety of surfactants to enable them to swarm across a semi-solid surface. For example, *P. syringae* pv. syringae B728a produces syringifactin and *P. aeruginosa* requires the presence of rhamnolipids to swarm (16–18). Swarming movement allows bacteria to move rapidly, search for and acquire food, avoid toxic substances, access and colonize new niches, and have increased resistance to antimicrobials (13, 19–22).

Motility as a requirement for bacterial root colonization and biocontrol has been shown for multiple bacterial species, but the majority of previous studies have focused on chemotaxis and swimming motility (23, 24). Comparatively fewer studies focus on swarming motility but work to date has shown a tight association between swarming, root colonization and biocontrol efficacy. For example, swarming defective but chemotactically proficient mutants (with intact swimming ability) of the PGPR *Bacillus subtilis* SWR01 have greatly reduced root colonization abilities (25). Similarly, a swarming mutant of *B. subtilis* 9407 could not colonize melon seedling roots and lost biocontrol efficacy against the pathogen that causes bacterial fruit blotch (26). Knockout studies of *P. parafulva* JBCS1880 showed that the coordinated action of antibacterial activity and swarming motility were critical for biocontrol against two soybean pustule-forming pathogens (27).

Knockout studies have long shown that flagella biosynthesis, assembly and regulatory genes are essential for swarming; mutants without flagella cannot swarm (28, 29). Studies are now showing that swarming involves more than just flagella genes (8, 14). A study using the *Escherichia coli* K12 Keio collection showed that knockouts of 7.4% of non-essential genes resulted in strongly repressed swarming (30). Yeung et al (31) showed that 233 *P. aeruginosa* PA14 transposon mutants had altered swarming motility, including flagellar or type IV pilus biosynthesis, transport, secretion, and metabolism genes. Knockout studies in *P. aeruginosa* PAO1 have shown that loss of polyphosphate kinase, encoded by the *ppk* gene and three genes relating to truncated lipopolysaccharide (*rmlC*, *migA* and *wapR*) result in a swarming deficient phenotype (13, 32). Swarming studies have also shown in *P. aeruginosa* PAO1 that *rsmA* in the Gac/Rsm regulatory pathway is a swarming regulator, and *bifA* phosphodiesterase mutants increase c-di-GMP levels and have reduced ability to swarm (33, 34).

Swarming motility is comparatively understudied in comparison to swimming motility, with studies mainly taking place in a mammalian infection context. Studies of swarming motility have uncovered substantial differences between swarming and other motility types. For example, swarming by pseudomonads requires more torque than other forms of motility, so during swarming these bacteria engage the MotC/MotD stator system (plus MotY) instead of the conventional MotA/MotB stator system of *E. coli* and *Salmonella* (35, 36). Studies of swarming motility in the pseudomonads have largely focused on regulation of swarming in the opportunistic human pathogen *P. aeruginosa* (37). Further research on swarming motility in species with different lifestyles and habitats will likely find additional differences in genes and proteins underlying swarming motility and its physiological and ecological importance. For example, swarming of *P. syringae* pv. syringae B728a, a phyllosphere colonizing bacterium, is thermoregulated (38), and the regulatory complexity of swarming in *P. fluorescens* Pf0-1, a soil bacterium which is naturally deficient in swarming, has only recently been uncovered (37).

*Pseudomonas protegens* Pf-5 (hereafter referred to as Pf-5) is a plant growth promoting bacterium that produces a range of antibacterial and antifungal compounds and can control pathogens of several crops (15, 39–43). The few swarming studies in Pf-5 have focused on the effect of the GacS/GacA two component regulatory system on swarming, including characterization of the *gacA* transcriptome of Pf-5 on swarming medium (44, 45). In Pf-5 a swarming deficient phenotype arises when either *orfA*, the orfamide A biosurfactant biosynthesis gene, *gacA* or *gacS* have mutations (44, 46). A lack of swarming motility in a Pf-5 flagella biosynthesis mutant *flhA* has been demonstrated multiple times (44, 45, 47). Pf-5 has also been observed to have reduced swarming motility on iron limited media (48).

Here we used the genome wide approach Transposon Directed Insertion-site Sequencing (TraDIS) to identify genes involved in Pf-5 swarming motility. TraDIS combines high-density random transposon mutagenesis and high throughput sequencing to rapidly study the fitness of all non-essential genes and link phenotype changes with specific genotypes (49–52). The data reported here sheds light on the suite of genes important for Pf-5 swarming motility beyond the well-known contribution of flagellar genes.

## Method

### Bacterial strains and media

*Pseudomonas protegens* Pf-5 was isolated from soil of a cotton field in Texas, USA (39) and a complete genome sequence has been generated for this organism (ENA accession number CP000076; 53). Pf-5 was routinely cultured using King’s Medium B (KMB; 2% w/v proteose peptone, 8.6 mM K2HPO4, 1.5% w/v glycerol, 6 mM MgSO4, pH 7.2; 54) at 27°C. Modified KMB agar plates for swarming and controls contained reduced proteose peptone (1% w/v), 5.28 mM K_2_HPO_4_, 6 mM MgSO_4_ and used Bacto^TM^ Agar (Becton, Dickinson and Company; 44).

### Transposon mutant library challenges

The role of genes in swarming motility was investigated using a previously constructed *Pseudomonas protegens* Pf-5 transposon mutant library (55). Briefly, a custom transposome was constructed using EZ-Tn*5* transposase (EpiCentre) and a transposon carrying a kanamycin (Km) resistance cassette isolated from the plasmid pUT_Km (56). The custom transposome was electroporated into freshly prepared electrocompetent Pf-5 cells allowing the transposon to randomly insert throughout the Pf-5 genome. A total of ∼500,000 colonies from four independent transformations were collected from LB-Km agar to create the saturated mutant library pool. Sequencing and analysis using the Bio-Tradis pipeline (50) indicated the library contains ∼256,000 unique transposon insertion sites spread evenly throughout the genome, with an average of one transposon insertion every ∼27 base pairs. This equates to an average of ∼45 transposon insertion sites per non-essential protein coding gene.

Five microliters of the Pf-5 transposon mutant library (5.87 x 10^7^ CFUs) was spotted on to the center of two swarm agar plates (modified KMB with 0.6% Bacto agar; 24 x 24 cm) in triplicate (two plates form each biological replicate with 1.17 x 10^8^ CFUs). After 24 h at 22°C, 4-5 mm of most motile cells were recovered from the edge of the swarming zone (Figure 1). The cells from the whole circumference of the swarming zone were scraped to the corner of the plate with a loop, washed off the agar surface with Phosphate Buffered Saline (PBS; Sigma Aldrich) and cells from two plates were collected to form each biological replicate (termed the output pool). For the controls 166 μL of the transposon mutant library (1.7 x 10^8^ CFUs) was spread on six modified KMB 1.5% agar plates (diameter 90 mm) in triplicate (total of 18 plates). After 24 h at 22°C the lawn was scraped off the agar surface of all six plates and collected in PBS (termed the control pool).

**Figure 1.**
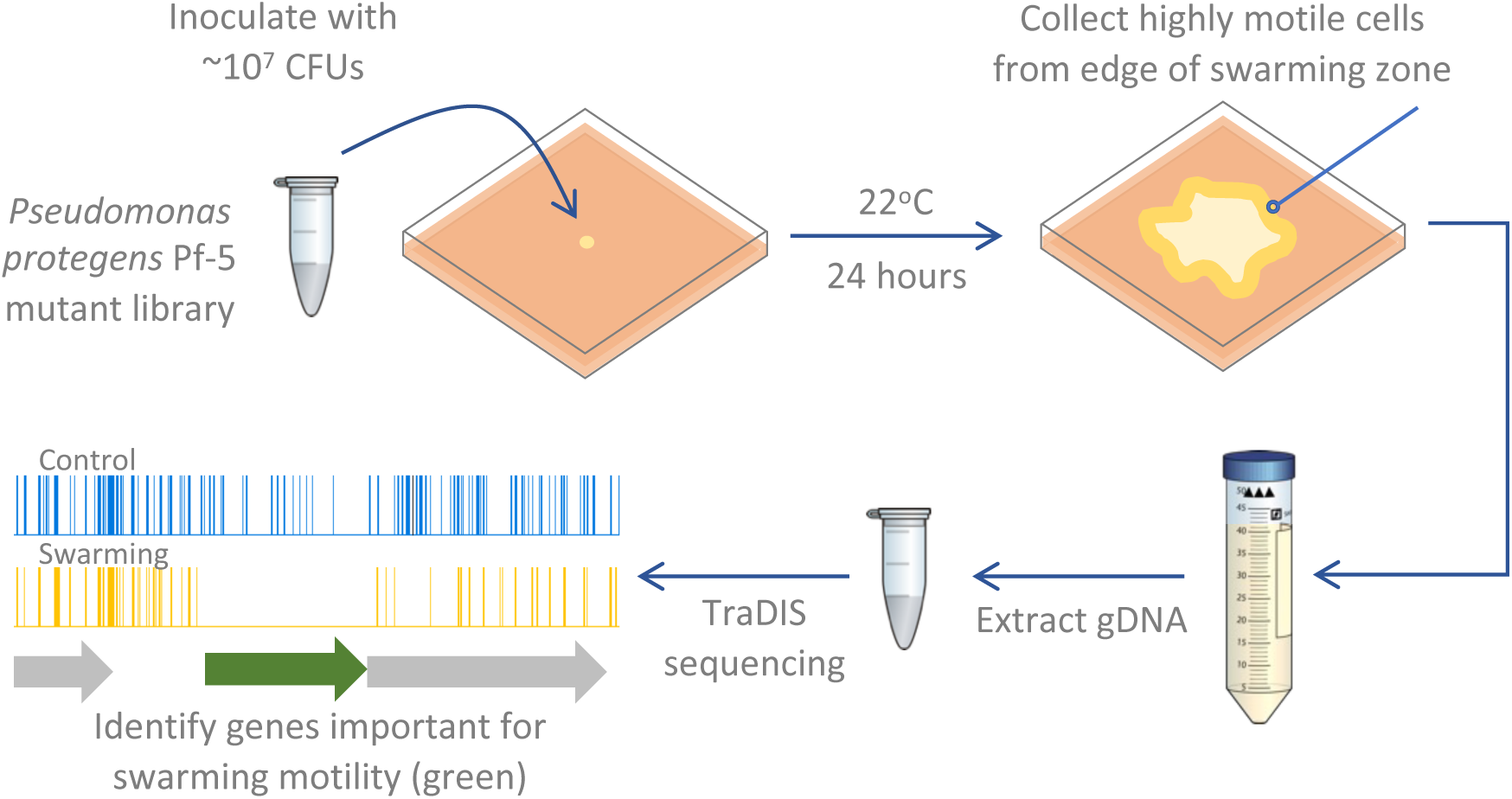
Schematic of the experimental methodology used to identify *Pseudomonas protegens* Pf-5 genes important for swarming motility. The example gene indicated in green in the last panel is important for swarming motility as the number of transposon insertion reads for this gene are significantly lower in the swarming assay compared to the control (reads visualized using Artemis software (57)).

### TraDIS sequencing and bioinformatic analysis

DNA was extracted from the cells from the output and control pools using a modified Wizard Genomic DNA Purification Kit protocol (Promega). The manufacturer’s protocol was followed with two modifications: centrifugation of the precipitated DNA was increased to 18,000 x g for 5 minutes and to 14,000 rpm for 10 minutes for the ethanol washed DNA. Contaminating agar was removed from the DNA using the Wizard SV Gel and PCR Clean-Up System (Promega) as per the manufacturer’s instructions. Transposon-directed insertion site sequencing (TraDIS) was performed at the Ramaciotti Centre for Genomics (UNSW, Sydney, Australia) in duplicate as previously described by Barquist et al (50) with an Illumina MiSeq platform to obtain 52 bp single-end genomic DNA reads. The transposon insertion sites were mapped to the Pf-5 genome and statistically analysed using the Bio-Tradis pipeline (50) as previously described (55). Briefly, this included allowing a 1bp mismatch in the transposon tag, excluding transposon insertions in the extreme 3’ end of each gene, and mapping reads with more than one mapping location to a random matching location. After matching the transposon tag, an average of 1.41 million reads per replicate were mapped to the Pf-5 genome (Table S1). A linear regression of the gene insertion indexes of the replicates was completed in R (58). Correlation coefficients between the insertion indexes for all pairs of replicates were >0.94 (p < 2.2 x 10^-16^; Figure S1) which validates the reproducibility of our replicates and is consistent with the reproducibility of transposon insertion sequencing replicates in other studies (59). The transposon insertion sites in each of the output pools were compared with those of the control pool with the Bio-Tradis tradis_comparison.R script with default parameters. Only genes with greater than 10 reads in both replicates of either the control or treatment condition were included to avoid genes being falsely classified as important for fitness. A cut-off of a log_2_ fold change of two in the number of transposon insertion reads in the output pools compared to the control pool and a q-value of < 0.01 was used to identify genes that when knocked out affected Pf-5 fitness during swarming.

Clusters of Orthologous Groups (COG) assignments (60) and Kyoto Encyclopedia of Genes and Genomes (KEGG) Orthology (KO) terms (61, 62) for each Pf-5 gene were previously compiled using eggNOG-mapper (63) with 87.3% of Pf-5 coding genes able to be assigned a COG code (55). The sum of all categories does not equal the total number of genes of interest, as some genes are assigned multiple COG codes, and some do not have a COG code assigned. Orthologs of Pf-5 genes were identified using the Pseudomonas Genome Database available at https://pseudomonas.com (64).

### Construction of knockout mutants

In-frame chromosomal gene deletion mutants were generated for Pf-5 genes PFL_4083, *pvdE* (PFL_4091) and PFL_5495 by creating gene deletion constructs via an overlap-extension PCR method, followed by allelic exchange with the suicide vector pEX18Tc (65), using a protocol adapted from Kidarsa et al (66). To create the mutant allele, fragments of 500-1100 bp flanking upstream and downstream of each gene of interest were first amplified by PCR from genomic DNA using the upstream (UpF/UpR) or downstream (DnF/DnR) primer pair. All PCRs were performed using KOD Hot Start DNA polymerase (Novagen) according to the protocol described by Kidarsa et al (66), using the primers listed in Table S2. The upstream forward primer and downstream reverse primer each had a 5’ extension adding an *Xba*I restriction site, while the upstream reverse and downstream forward primers were designed to be in-frame with the gene of interest and had a 5’ linker of 12 bp complementary to each other to allow overlapped annealing during the secondary PCR. The amplicons of the upstream and downstream primary PCRs were gel-purified, mixed together 1:1 (50 ng each) and used as the template for the secondary PCR using the UpF and DnR primers. The resultant full-length product was gel-purified, digested with *Xba*I, treated with Calf Intestinal Alkaline Phosphatase (New England Biolabs), and then cloned into the pEX18Tc vector (linearised with *Xba*I) using T4 ligase (New England Biolabs).

The recombinant vectors were transformed by electroporation into ElectroMAX DH5α-E competent *E. coli* cells (ThermoFisher Scientific) and the mutant alleles were verified by PCR and sequencing using pEx18Tc sequencing primers (Table S3). All vectors were subsequently electro-transformed into competent cells of the mobilising strain *E. coli* S17-1 (67).

Biparental matings were performed between the vector-containing *E. coli* S17-1 and wild-type Pf-5 for conjugative transfer of each vector into Pf-5 as described by Lim et al (68) but using nutrient agar containing 1.5% (v/v) glycerol. Transconjugant Pf-5 colonies were selected on KMB agar containing 200 µg mL^-1^ tetracycline (vector conferred resistance) and 100 µg mL^-1^ streptomycin (innate resistance of Pf-5). The surviving colonies were grown without selection in LB broth for 3 h with shaking and plated on LB with 1.5% agar supplemented with 10% sucrose to resolve merodiploids (due to counter-selection against *sacB*-carrying cells). Sucrose-resistant colonies were patched in parallel onto LB with 1.5% agar containing 10% sucrose and KMB agar with 200 µg mL^-1^ tetracycline to further confirm the absence of the pEX18Tc vector backbone. Tetracycline-sensitive colonies were screened by PCR with primers annealing to chromosomal regions external to the target gene (Table S3) to detect mutants with truncated amplicon sizes compared to a wild-type control. The deletion of each gene was confirmed by PCR of genomic DNA from each Pf-5 mutant colony with subsequent sequencing before being stored in 25% glycerol at −80°C until required.

Confirmed deletion mutants were grown on half strength litmus milk agar (Fluka Analytical) at 27°C overnight to ensure they did not have spontaneous g*acA* or *gacS* mutations. A cleared zone, or halo, around bacterial colonies indicated there was extracellular protease activity and there were no *gacA* or *gacS* mutations present (69, 70).

Growth curves were conducted to check for general growth defects in knockout mutants. Overnight cultures of wildtype Pf-5 and the mutant strains ΔPFL_4083, ΔPFL_4091, and ΔPFL_5495 were each grown at 27°C with shaking for 16 hours in Mueller-Hinton II (MH) broth. The cultures were sub-cultured 1:25 into fresh MH broth and incubated with shaking until OD_600_ = 0.6. Cultures were kept on ice while serial dilutions were performed in MH broth to reach a final density of 1.2 x 10^4^ CFU/mL in 150 µl MH broth in a 96-well plate (10 replicates per strain). Plates were incubated in a Pherastar plate reader at 27°C with shaking at 200 rpm and OD_600_ readings were taken at 6-minute intervals for 24 hours.

### Phenotypic assays with knockout mutants

Motility assays were performed on modified KMB soft agar plates, freshly prepared as above and allowed to set for at least 4 hr at 22°C prior to inoculation. To prepare the inoculum, cells of Pf-5 wild-type and each mutant were scraped from a culture grown on KMB agar for 48 h, then resuspended in PBS and diluted to an OD_600_ of 0.2. Two microlitres of the cell suspension was spotted onto the surface of modified KMB soft agar plates in quadruplicate. All cultures were incubated at 22°C for 24 hr and the diameter of the motile zones were measured. One-way ANOVA followed by post-hoc Tukey’s HSD test was conducted to compare the swarming diameter of each mutant to the parental strain.

A droplet collapse assay was used to test for surfactant activity of the Pf-5 knockout mutants (71). Bacterial cultures grown for 48 hr at 27°C on KMB agar plates were suspended in PBS (Sigma Aldrich) to a final density of 1 x 10^10^ CFU mL^-1^ and 10 μl droplets were spotted on parafilm. Three replicate drops were used for each culture. A flat droplet indicated the cells produced the surfactant orfamide A, while domed droplets indicated the cells did not produce orfamide A (45).

The Pf-5 mutants were streaked onto KMB 1.5% agar plates and incubated at 27°C for 48 hours. The coloration of the cells was observed under fluorescent lights (white light) and the fluorescence of the streak plates was observed using UV excitation.

## Results and Discussion

### Identifying genes contributing to Pf-5 swarming fitness

The swarming experiments using the *Pseudomonas protegens* Pf-5 saturated transposon mutant library identified 136 genes where loss of function affected Pf-5 swarming fitness (2.40% of the non-essential genes; 55; Figure 2). Loss of function of 119 of these genes was detrimental for Pf-5 swarming fitness, based on cells carrying mutations in these genes being present at significantly lower level in the output pool than the control pool (Figure 2). A set of 21 genes were identified where loss of function led to enhanced Pf-5 swarming fitness, with cells with mutations in these genes present at significantly higher levels in the output pool than in the control pool (Figure 2). The fold changes for the full list of Pf-5 genes are available in Dataset S1.

**Figure 2.**
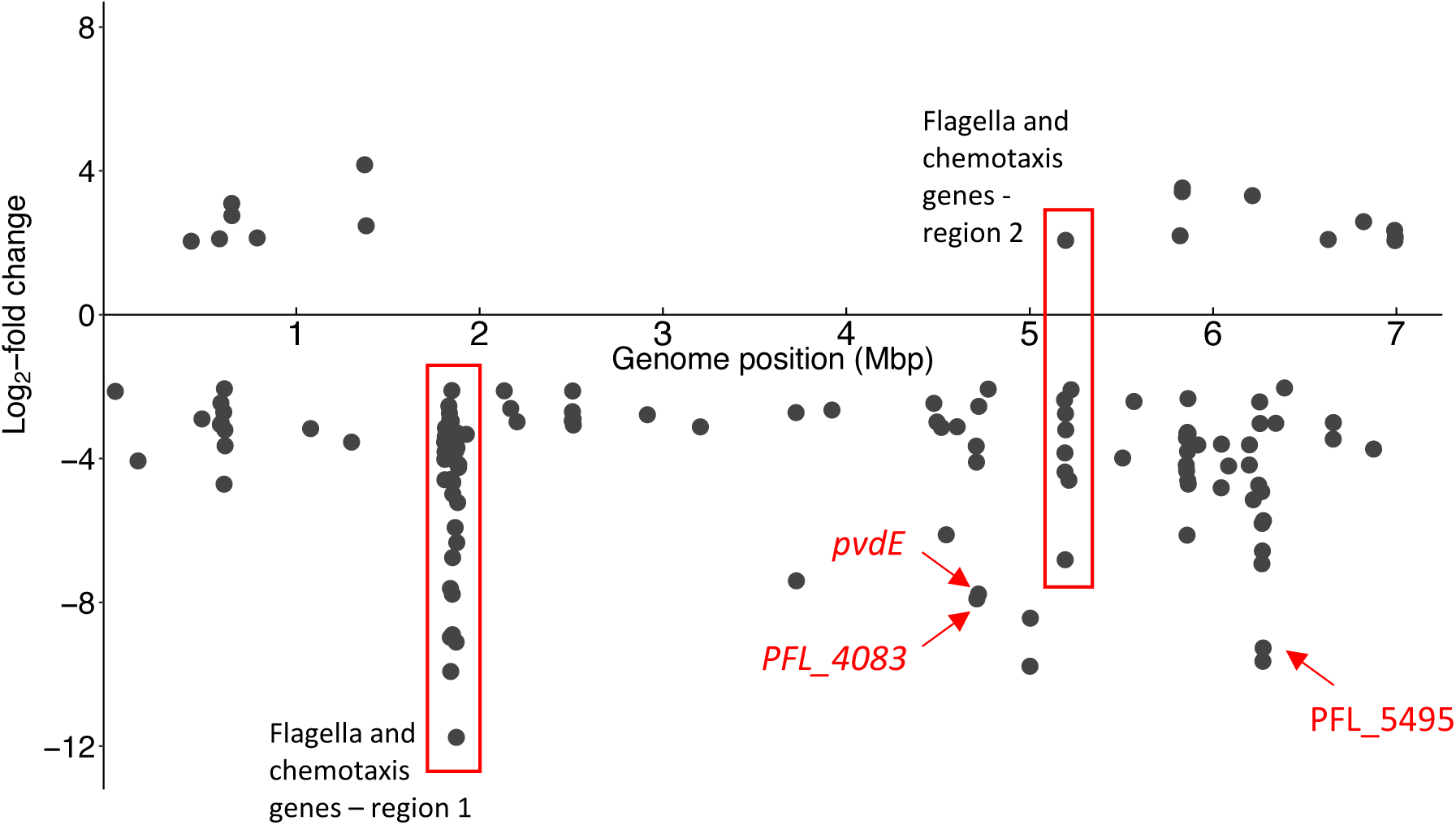
Genome position of *Pseudomonas protegens* Pf-5 genes identified by TraDIS with log_2_ fold change > 2 or < −2 from swarming assays when compared with the control. Genes with a log_2_ fold change of less than −2 indicate that loss of their function was associated with reduced swarming fitness and greater than 2 indicate that loss of their function was associated with enhanced swarming fitness. Genes that were knocked out in this study are indicated with red text. Scatterplot generated using the R package ggplot2 (72).

### Functional analysis of genes associated with swarming fitness

A functional analysis of the Pf-5 genes that affected swarming fitness was obtained by classifying the genes using Clusters of Orthologous Groups (COG) categories (60). The functional category of cell motility (N) made up the largest group of genes that detrimentally affected swarming fitness when their function was lost (Figure 3). Other categories which contained substantial numbers of genes for which loss of function reduced swarming fitness were cell wall/membrane/envelope biogenesis (M), signal transduction mechanisms (T), coenzyme transport and metabolism (H) and unknown function (S; Figure 3). There were only 21 genes for which loss of function was associated with enhanced swarming fitness, the largest proportion of which are in the functional category of Transcription (K; Figure 3).

**Figure 3.**
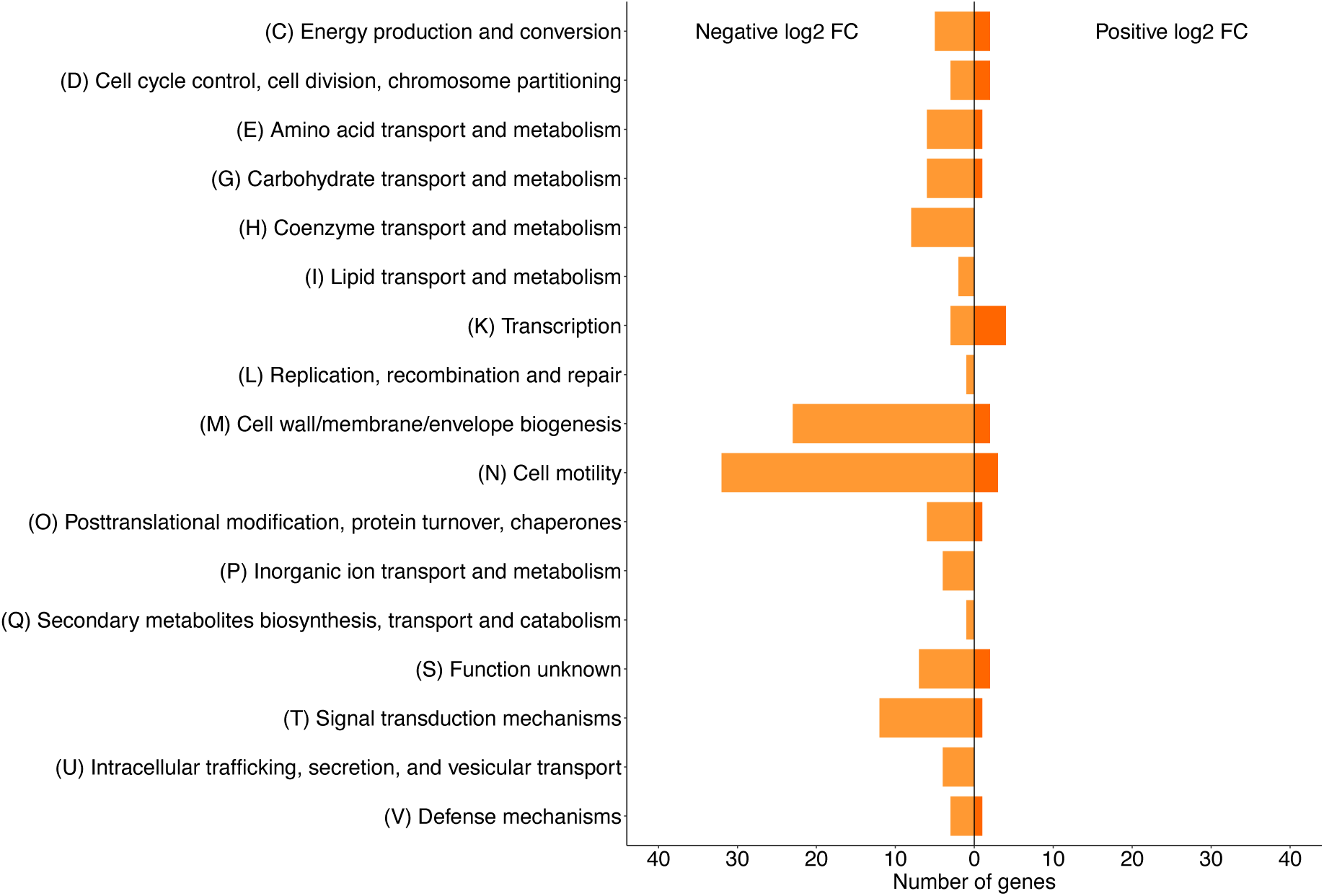
Functional classification of genes that affect swarming fitness when their function is lost (log_2_ fold change > 2 or < −2). Data visualized using the R package ggplot2 (72).

### Swarming motility requires many flagellar genes

Flagella are known to be essential for swarming motility (28, 29). There are 48 genes related to flagella structure, biosynthesis, assembly, and regulation in the Pf-5 genome located in two major clusters (73). Our TraDIS data showed that loss of function of 37 of these genes affected Pf-5 swarming motility (Figure 4). For 34 of these flagellar genes loss of function reduced swarming fitness, which is consistent with previous data for Pf-5 and other pseudomonads (Figure 4; 31, 44, 47, 74, 75). In contrast, loss of function of three genes, *motAB* and PFL_4484, enhanced Pf-5 swarming fitness. MotAB and MotCD are the two sets of stators in the flagellar motor that generate torque (76). Either set of stators are sufficient for swimming, but swarming requires the MotCD stator system (35, 77). In *P. aeruginosa* PA14 swarming motility increased when *motAB* were knocked out (78), which is consistent with the enhancement of Pf-5 swarming fitness when the functions of *motAB* are lost.

**Figure 4.**
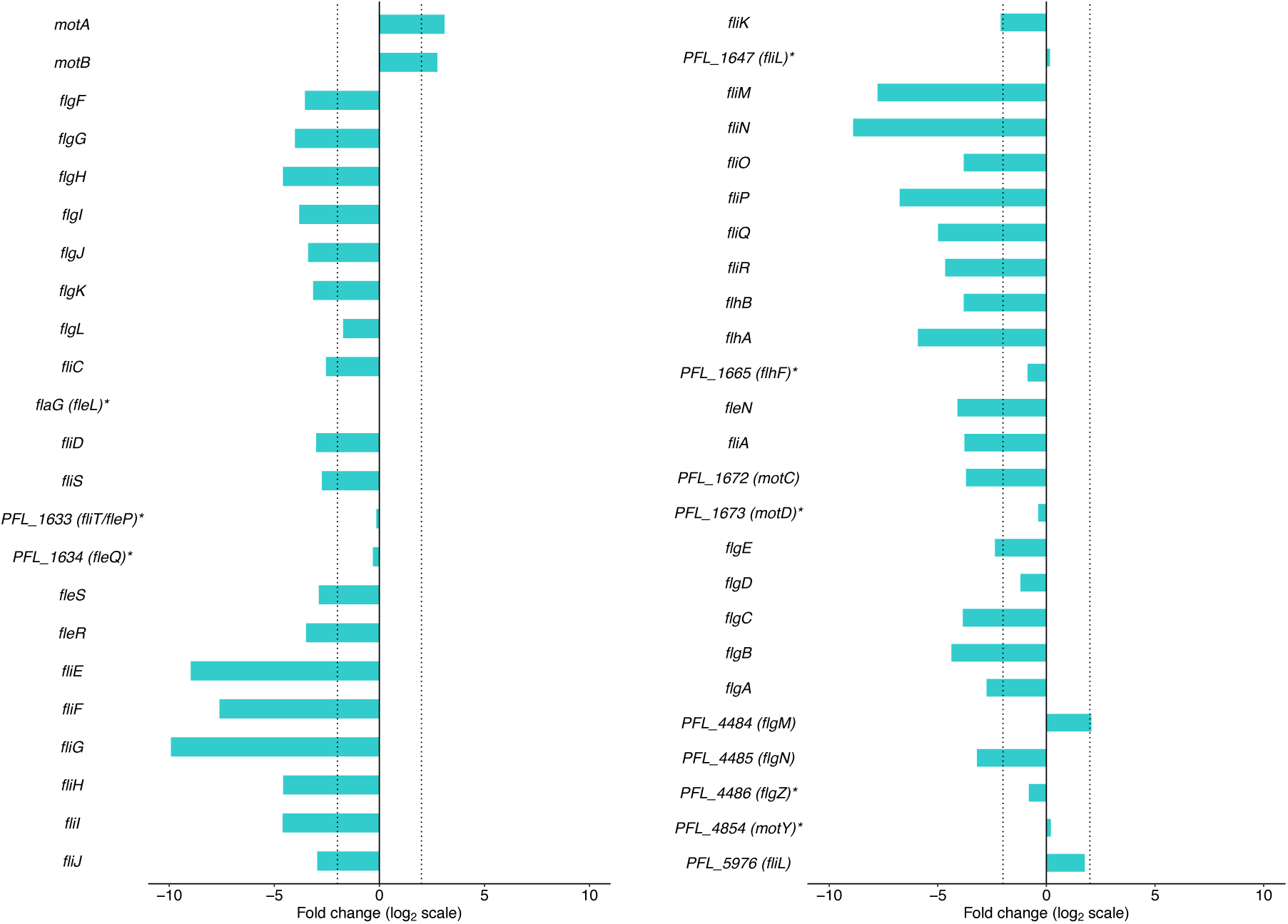
Log_2_ fold change in read abundance of *Pseudomonas protegens* Pf-5 flagella-related genes during swarming. Dotted vertical lines are positioned at log_2_ fold changes of −2 and 2. Orthologous gene names from *P. aeruginosa* are in parentheses. *Genes with non-significant log_2_ fold change. Data visualized using the R package ggplot2 (72).

Mutation of the regulator gene PFL_4484 that encodes the anti-sigma-28 factor FlgM also enhanced Pf-5 swimming motility. During flagellar synthesis in *P. aeruginosa*, FlgM negatively regulates flagella gene expression by binding sigma 28 factor FliA (79). Loss of FlgM function allows FliA regulated genes to be transcribed in the same way as when FlgM is prevented from binding FliA by the anti-anti-sigma factor HsbA (80).

### Cell envelope biogenesis and cell division

There were 36 genes relating to cell division and cell wall/membrane biogenesis that affected Pf-5 swarming motility when their function was lost (Figure 5A). Twenty-eight of these genes relate to lipopolysaccharide (LPS) biosynthesis, three to peptidoglycan turnover, and five to cell division. Loss of 32 of these genes detrimentally affected swarming, with ten having a log_2_ FC < −5 indicating loss of these genes had a very strong negative effect on swarming.

**Figure 5.**
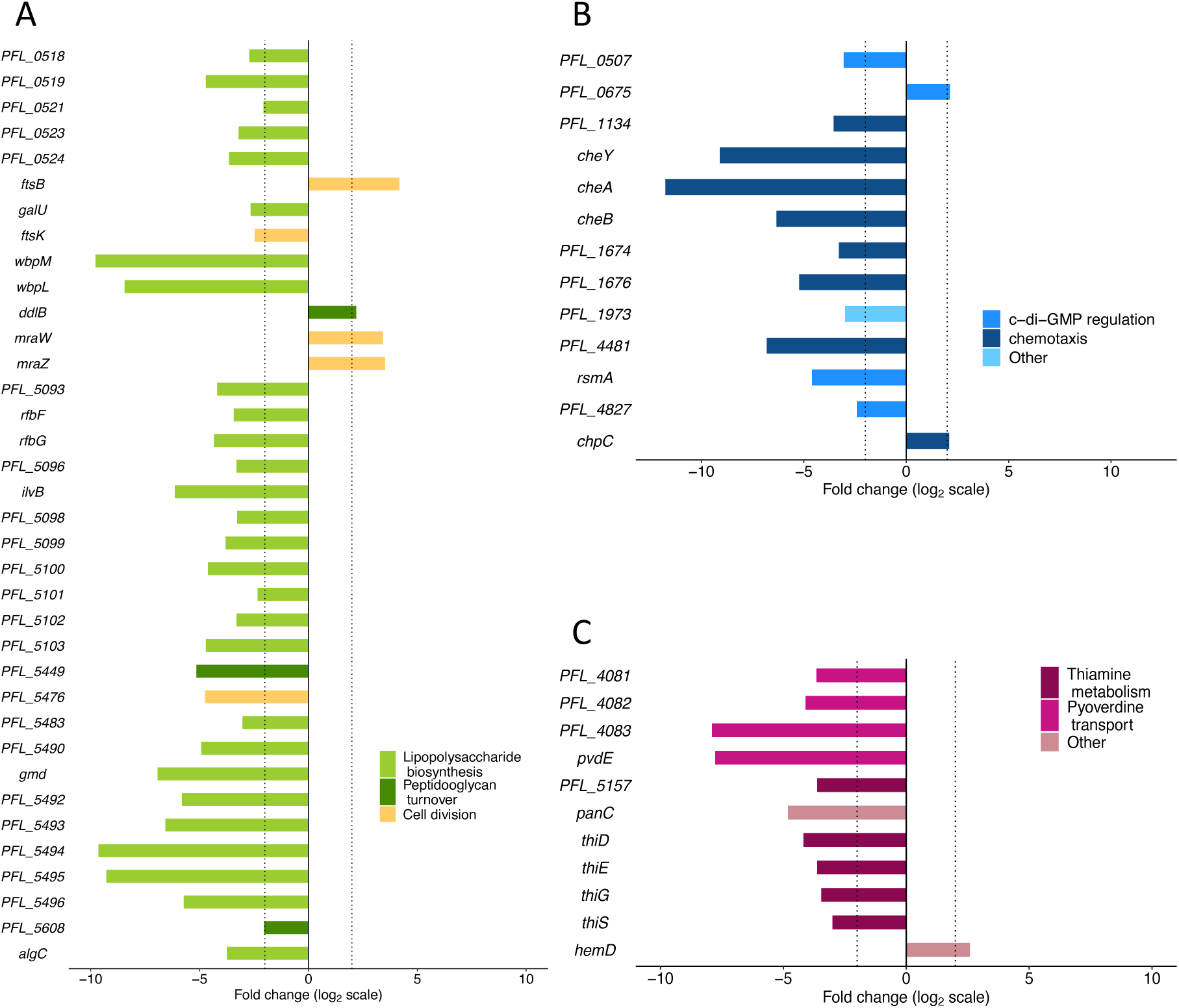
Log_2_ fold change in read abundance of *Pseudomonas protegens* Pf-5 genes related to (A) cell envelope biogenesis and cell division, (B) signal transduction, and (C) coenzyme transport and metabolism during swarming. Dotted vertical lines are positioned at log_2_ fold changes of −2 and 2. Data visualized using the R package ggplot2 (72).

Twenty-eight of the cell wall/membrane biogenesis genes showing effects on Pf-5 swarming motility have functions in LPS biosynthesis, including genes associated with LPS core biosynthesis (PFL_0518, PFL_0521, PFL_0524, *galU*, *algC*) as well as those related to O-antigen biosynthesis (wbpM, wbpL, rfbF, rfbG, rfbH, rfbV, PFL_5483; 81) and common polysaccharide antigen (PFL_0519, PFL_5490, gmd, PFL_5494, PFL_5495; 82). Loss of all these genes detrimentally affected swarming and nine had a very strong negative effect (Figure 5A). One operon, containing two glycosyltransferases and two components of an ABC transporter, had all four genes showing a strong negative fold change (PFL_5492– PFL_5495; Figure 5A). Mutants of genes related to LPS biosynthesis have been shown to have absent or truncated *O*-antigens or core LPS. For example, *wbpL* mutant strains of *P. syringae* pv. tomato DC3000 and *P. cichorii* ATCC10857/DSM50259 lacked *O*-polysaccharide (83) and *P. aeruginosa* serogroup O11, O5, O6 and O17 strains with mutated *galU* had truncated core LPS (84). Bacterial strains with truncated LPS often have motility defects compared to those with wild type LPS (32). LPS functions as a surfactant which reduces surface tension and decreases resistance between the cell and the surface (30). A genome wide screen of *E. coli* K12 mutants found that genes involved in LPS biosynthesis are required for swarming (30). A similar phenomenon has been reported in other Gram-negative bacteria, such as *S. enterica* serovar Typhimurium, *Proteus mirabilis*, *P. syringae* pv. tomato DC3000 and *P. cichorii* ATCC10857/DSM50259 (83, 85–87). In *P. aeruginosa* PAO1, motility defects in mutants with truncated LPS were due to increases in adhesion forces between the cells and cell to surface (32). The expression of flagellar genes is also impaired by cell envelope stress, specifically the truncation of LPS in *E. coli* and *S. enterica* serovar Typhimurium (88). These findings are consistent with the loss of 28 Pf-5 genes related to LPS biosynthesis resulting in impaired swarming motility.

Defects in the function of peptidoglycan-related proteins can result in abnormal cell shapes with non-standard cell aspect ratios (89). PFL_5449, observed here to be important for swarming, is an ortholog of *rlpA* in *P. aeruginosa* PAO1. RlpA is involved in degradation of peptidoglycan in the cell division septum and lateral cell walls, so it affects maintenance of rod shape (90). Abnormal shape (cell aspect ratio) interfering with swarming cell dynamics is observed in *Bacillus subtilis* 3610; cells that are different lengths to wild type cells have lower motility (91). This is consistent with the loss of PFL_5449 having a strong negative effect on swarming motility (Figure 5A).

Similarly, loss of genes with cell division functions can affect cell shape (92). One cell division-related gene, here implicated in Pf-5 swarming, is *ftsK* which encodes a DNA translocase involved in cell separation (cytokinesis) and chromosome partitioning (93). In *E. coli*, mutants with loss of function of *ftsK* have impaired cell-cell separation, resulting in chains of cells (94). This alteration in cell morphology with the loss of *ftsK* is consistent with detrimental affects on swarming motility with loss of this gene in Pf-5 and *P. aeruginosa* PA14 (31).

In contrast, loss of cell division-related genes *ftsB, mraW* and *mraZ* enhance Pf-5 swarming motility (Figure 5A). FtsB forms part of the FtsQLB complex that recruits and regulates the essential peptidoglycan synthase (FtsWI) responsible for septal peptigoglycan synthesis during cell division (95–97). In *P. aeruginosa* PAO1 the FtsQLB complex acts as an activator of FtsWI (98–100). Considering the negative impact of defects in peptidoglycan synthesis on cell division, cell shape and motility, it is not clear why loss of *ftsB* function enhances Pf-5 swarming motility.

Genes *mraW* and *mraZ*, located in the same operon as *ftsI*, also enhanced Pf-5 swarming motility when their function was lost. The gene *mraW* encodes MraW, a ribosomal RNA methyltransferase (101). In *E. coli* O157:H7, *mraW* mutants had genome wide decreased methylation, including flagella-related genes, resulting in reduced flagella production (102). Given that swarming motility relies on flagellar gene function, it is unclear why loss of function of *mraW* resulted in enhanced swarming motility of Pf-5. MraZ, a DNA-binding transcription factor encoded by *mraZ*, regulates its own expression as well as that of other cell division and cell wall genes (103). In *E. coli*, *Bacillus subtilis* and *Mycoplasma genitalium* MraZ acts as a transcriptional inhibitor of cell division, but in *M. gallisepticum* MraZ is an activator (103–105). The mechanism by which loss of *mraZ* function affects Pf-5 swarming motility is unknown.

### Signal transduction mechanisms

Loss of function of 13 signal transduction genes affected Pf-5 swarming motility; 11 of these genes were detrimental for swarming motility and 2 genes enhanced swarming motility (Figure 5B). Six of these genes are part of the Che chemotaxis pathway, with five having a strong detrimental effect on Pf-5 swarming motility (log_2_ fold change < −5). In *P. aeruginosa* PA14, some chemotaxis genes have been shown to be important for swarming, but the mechanism does not appear to be via changes in chemotactic behavior (9, 106). Similarly, in other bacteria the mechanism by which chemotaxis genes play a role in swarming motility is not yet elucidated, but these genes are central for motility with some aspects of the chemotaxis signal transduction phosphorelay essential for swarming cell differentiation (12, 107). This is consistent with reduced Pf-5 swarming motility resulting from the loss of function of chemotaxis genes.

Three of the signal transduction genes important for Pf-5 swarming are orthologs of genes in the c-di-GMP regulation system in pseudomonads (PFL_0507, PFL_0675, PFL_4827; Figure 5B). A variety of cellular processes are controlled by turnover of c-di-GMP, including motility, and a low intracellular concentration of c-di-GMP is associated with motile cells (108). Orthologs of PFL_4827 (*bifA*) and PFL_0507 (*dipA*) encode phosphodiesterases that breakdown c-di-GMP and when knocked out cause accumulation of c-di-GMP in cells and reduced motility. Knockouts of *bifA* and *dipA* have been shown to reduce motility in *P. aeruginosa* PA14 and PAO1, respectively, consistent with the detrimental effects of disrupting their orthologs on Pf-5 swarming motility (34, 109). Loss of function of PFL_0675, encoding a diguanylate cyclase protein, enhanced Pf-5 swarming motility (Figure 5B). Reduced synthesis of c-di-GMP due to the disruption of diguanylate cyclases has been reported to enhance swarming motility as intracellular c-di-GMP levels are lower (108). Loss of the *wspF* ortholog PFL_1134 from the Wsp pathway also detrimentally affected Pf-5 swarming motility (Figure 5B). The absence of the methylesterase WspF in *P. aeruginosa* PA14 causes constitutively high levels of c-di-GMP levels through continuous activation of the response regulator WspR and a hyper-biofilm phenotype (110). This is consistent with a detrimental effect on Pf-5 swarming motility when the *wspF* ortholog PFL_1134 is lost.

Loss of *rsmA* was detrimental for Pf-5 swarming motility (Figure 5B). Knocking out *rsmA* in *P. aeruginosa* PAO1 and PA14 causes a loss of swarming motility (31, 33). RsmA is a small RNA-binding regulatory protein, a global regulator in the GacA/GacS two component system, that represses translation of its target genes, one of which is *algU* (111). In *P. fluorescens* F113 and *P. aeruginosa* PAO1 the loss of function of *rsmA* results in the loss of repression of *algU*, which encodes an activator of *amrZ* transcription (112, 113). AmrZ is a transcriptional regulator of *fleQ*, the gene encoding the master flagellar regulator in both these strains (111, 114). Flagella biosynthesis and function are essential for swarming, so the negative effect on flagella biosynthesis from this cascade resulting in *fleQ* repression is consistent with the loss of *rsmA* being detrimental for Pf-5 swarming motility.

### Co-enzyme transport and metabolism

Seven genes relating to co-enzyme transport and metabolism affected Pf-5 swarming motility when their function was lost: six affected swarming detrimentally and one enhanced swarming motility (Figure 5C). Five of the genes important for swarming motility are in the thiamine (vitamin B_1_) biosynthesis pathway (115). To our knowledge, a link between thiamine biosynthesis and swarming motility has not been previously reported.

Thiamine, and its active form thiamine pyrophosphate, is crucial for bacterial survival as it is an essential co-factor for several carbon metabolism enzymes, including those involved in glycolysis, the tricarboxylic acid (TCA) cycle, and the pentose phosphate pathway (115–117). For example, thiamine is essential for colonization and survival of *P. syringae* pv. tomato DV3000 and *P. fluorescens* WCS365 on tomato plants (118, 119). Knocking out a thiamine transporter operon (*thiBPQ*) in *Edwardsiella piscicida* ZW1 correlated with a reduced intracellular c-di-GMP level (120). This suggests that thiamine biosynthesis might affect swarming motility via c-di-GMP levels. Further studies into this may shed more light on the connection between this essential enzyme co-factor and swarming motility.

### Swarming motility requires conserved genes of unknown function

Loss of 4 genes encoding hypothetical proteins affected Pf-5 swarming motility, with three of these reducing Pf-5 swarming fitness. One of these genes, PFL_5098, is likely related to cell envelope production. It is part of a cluster of 11 O-antigen biosynthesis genes that all have detrimental effects on swarming fitness when their function is lost (PFL_5093– PFL_5103). One of the genes encoding a hypothetical protein, PFL_0130, had a log_2_ fold change of −4.07, indicating that it had a strong negative effect on swarming fitness and making its characterization of particular interest as it may provide further insights into swarming motility in this plant-associated bacteria.

### Investigating efflux transporters important for swarming motility

Three single knockout mutants of Pf-5 (Δ*pvdE*, ΔPFL_4083 and ΔPFL_5495) were created for experimental validation and further investigation as they showed a large negative fold change in reads in swarming assays (Figure 2). All three genes encode parts of efflux systems associated with pyoverdine (*pvdE* and PFL_4083) or lipopolysaccharide (PFL_5495) transport. There was a significant difference in swarming diameter of the mutants when compared to the parental strain (ANOVA, F(3,51) = 22.8, p < 0.01). All the mutants had a smaller swarming diameter than the parental strain, with the swarming diameters of Δ*pvdE* and ΔPFL_4083 the smallest (post-hoc Tukey’s HSD; Figure 6; representative images in Figure S4). These results confirm our TraDIS findings and suggest that TraDIS has accurately identified a suite of genes involved in Pf-5 swarming motility. Growth curves for all single knockout mutants confirmed that there were no generalized growth defects present (Figure S3). In a droplet collapse assay all mutants had flat droplets which indicates that production of the biosurfactant orfamide A was normal (45). Litmus milk assays with each mutant had a cleared zone around the inoculation point which confirmed exoprotease production was normal and there were no secondary *gac* mutations present (70, 121).

**Figure 6.**
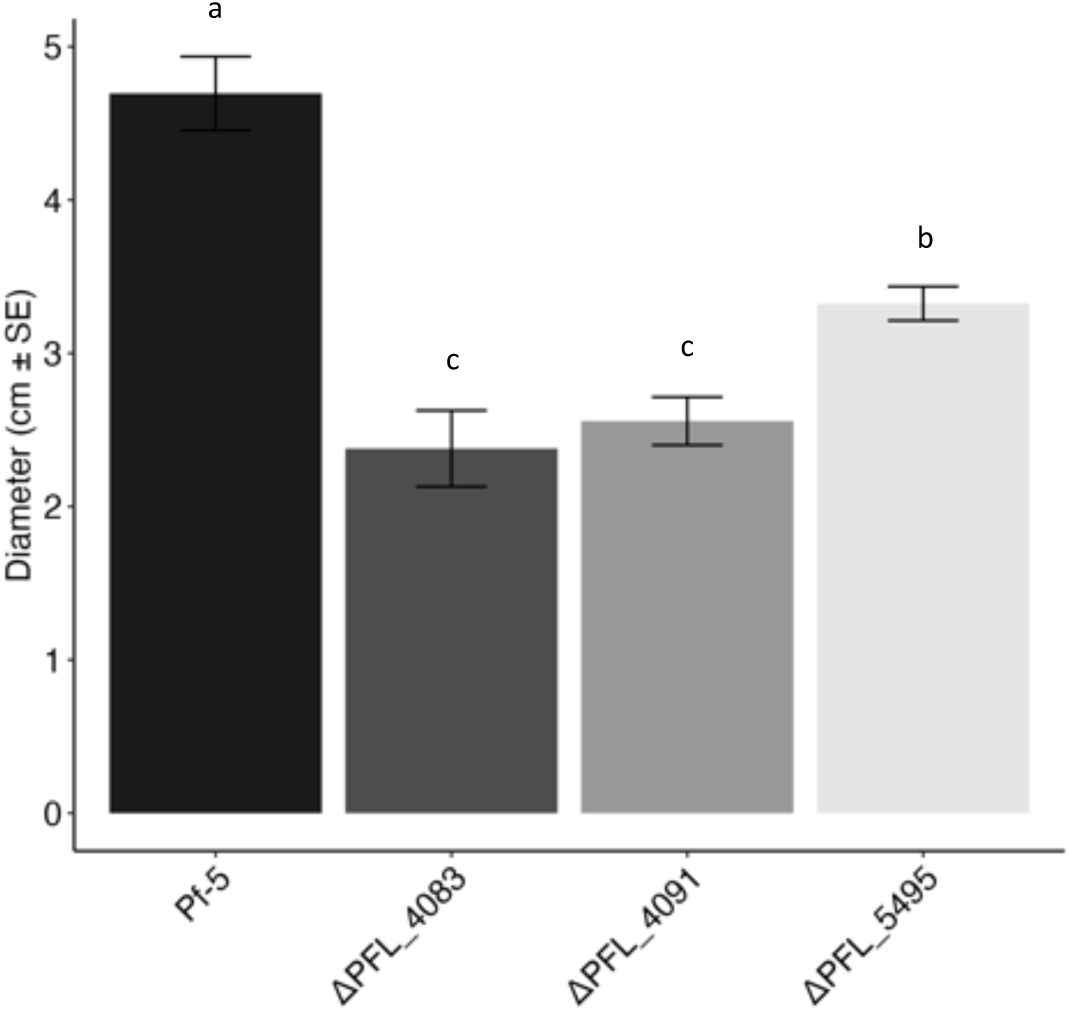
(A) Swarming assays conducted with *Pseudomonas protegens* Pf-5 single knockout mutants. Mean diameter (cm ± SE) of the motile zones was measured after 24 hours growth at 22°C on modified KMB soft agar plates (0.6% agar). Letters indicate significant difference at a = 0.05 determined using ANOVA and post-hoc Tukey’s HSD Test. Figure generated using the R package ggplot2 (72).

PFL_4083 and *pvdE* are part of the Pf-5 pyoverdine biosynthesis gene cluster (PFL_4077-4140, PFL_4146-4160, PFL_4162-4191; 53). Pyoverdine is a fluorescent siderophore produced by pseudomonads for iron acquisition from the environment (122). The bacteria produce pyoverdine in the cytoplasm, mature it in the periplasm and secrete it into the extracellular medium (123). The siderophore binds to extracellular ferric iron and the complex is transported back to the bacterial periplasm via an outer membrane transporter (124). The iron is released from the complex in the periplasm and the pyoverdine is re-used (125, 126). Siderophores are traditionally considered ‘public goods’ in that secreted siderophores are available to all cells in the population, including those not producing siderophores, known as ‘cheaters’, which can take up siderophores produced by others and obtain an advantage (127). Under this scenario, genes related to the production of public goods, for example pyoverdine-related genes, would not be expected to be identified using TraDIS (128) as the transposon insertion library is a mixture of cells including those without the capacity to produce pyoverdine (disruption of pyoverdine biosynthesis, transport or recycling genes) and cells that can produce pyoverdine. Studies of pyoverdine uptake in a mixed community of producers and non-producers of *P. aeruginosa* PAO1 have shown that during growth on a solid surface pyoverdine is localized and not widely diffused as in liquid culture, so non-producers may not have access and therefore may not benefit as much as previously thought (129).

PvdE is an ABC transporter which translocates the pyoverdine precursor ferribactin from the cytoplasm to the periplasm where is undergoes maturation to the fluorescent pyoverdine (130). Observation under white and UV light showed that Pf-5 Δ*pvdE* was deficient in pyoverdine while pyoverdine was present in the parental strain (Figure S5). The *pvdE* knockout also had smaller colonies on KMB agar plates (a low iron media commonly used for growth of fluorescent pseudomonads), likely due to limited growth due to no pyoverdine production (48). The PvdE efflux pump has been shown to have a critical role in pyoverdine production in *P. aeruginosa* PAO1 (126) and Pf-5 pyoverdine biosynthesis genes are upregulated in response to iron limitation (48). Disruption of the biosynthesis and transport of siderophores in *P. aeruginosa* PA14 has been linked to reduced swarming motility (131, 132). Similarly, pyoverdine mutants of *P. putida* KT2440 lose the ability to swarm (133). To our knowledge there are no studies directly knocking out *pvdE* in a pseudomonad, but *pvdE* has been shown to be upregulated during *P. aeruginosa* PA01 swarming (134). Pf-5 has reduced swarming motility under iron limited conditions (48), consistent with the detriment to Pf-5 swarming fitness when the function of *pvdE* is lost.

Genes from the operon PFL_4081-PFL_4083 are orthologs of the genes *pvdR*, *pvdT* and *ompQ*, respectively, that encode the three components of the efflux pump PvdRT-OmpQ involved in secretion of newly synthesised and recycled pyoverdine from the periplasm (123, 126, 135). The contribution of the efflux pump PvdRT-OmpQ to pyoverdine secretion may vary among pseudomonads. For example, in *P. aeruginosa* PAO1, knocking out *ompQ* results in an increase in pyoverdine in the periplasm, but has no effect on secretion of newly synthesized pyoverdine (123). However, in *P. taiwanensis*, a soil biocontrol bacterium, the PvdRT-OmpQ system is not involved in pyoverdine secretion (136). The PFL_4083 knockout grown in low iron conditions produced and secreted fluorescent pyoverdine and had similar colony size to the parental strain (Figure S5). Further study of the Pf-5 PvdRT-OmpQ complex is needed to determine which aspects of pyoverdine secretion this transporter conducts in this species. There are no motility studies of *ompQ* knockouts to our knowledge, but mutants of *pvdR* in *P. aeruginosa* PA14 had significantly impaired swarming (131). Loss of function of the *P. aeruginosa* PA14 PvdRT-OmpQ complex reducing swarming motility is consistent with the loss of these orthologous genes detrimentally affecting Pf-5 swarming motility.

PFL_5495 is part of the operon PFL_5492-PFL_5495 which includes genes encoding two glycosyltransferases and two components of an ABC transporter. Loss of function of any of the four genes from this operon had a strong negative effect on Pf-5 swarming motility (log_2_ fold change < −5; Figure 5A). ΔPFL_5495 had reduced swarming compared to the parental strain (Figure 6). PFL_5495 is an ortholog of *wzm* from *P. aeruginosa* PAO1 which encodes the membrane subunit of the A-band LPS efflux transporter. PFL_5494 from the same operon is an ortholog of *wzt* from *P. aeruginosa* PAO1, encoding the ABC-subunit of the A-band LPS efflux transporter.LPS is made up of lipid A, core oligosaccharide and O-polysaccharide (also referred to as O-antigen). There are two types of O-polysaccharide: A-band, also called Common Polysaccharide Antigen, and B-band, also termed O-Specific Antigens (137). In *P. aeruginosa* strains, Common O-Polysaccharide chains are fully assembled on the cytoplasmic side of the inner membrane and then exported by an ABC transporter to the periplasmic face (137). The genes *wzt* and *wzm* encode the two components of the ABC O-polysaccharide transporter in *P. aeruginosa* PAO1. Mutation of either of these genes prevents export of Common O-Polysaccharide (A-band) to the periplasm and ultimately to the cell surface (138). Detrimental effects of truncated LPS and absent O-polysaccharide on swarming motility are discussed above. For example, swarming of *P. putida* KT2440 requires LPS O-antigen (133), consistent with the detrimental effect on Pf-5 swarming motility when PFL_5495 function is lost.

### Conclusions

In this study 136 genes involved in swarming motility of *P. protegens* Pf-5 were successfully identified using TraDIS. This genome-wide method identified genes that are known to contribute to swarming as well as genes not previously associated with this type of motility. Successful swarming required the function of genes related to flagella biosynthesis, assembly, function and regulation, signal transduction genes including chemotaxis (but not chemotactic sensing) and c-di-GMP turnover. We also identified genes for which the loss of function detrimentally affected swarming motility, including cell division, peptidoglycan turnover and LPS biosynthesis genes, with their loss likely affecting cell morphology and surface adhesion properties. Loss of function of genes involved with the metabolism of thiamine (vitamin B_1_) and transport of pyoverdine also had a large negative effect on swarming motility.

This study expands our knowledge of swarming motility, both in the breadth of genes identified and by examining a species with a different lifestyle and habitat than the majority of existing studies. Motility, root colonization and biocontrol function are tightly linked, so increasing our understanding of the genetic basis of swarming motility enhances our knowledge of key processes of plant growth promoting rhizobacteria required for their efficacy in agricultural settings.

## Acknowledgements

*Pseudomonas protegens* Pf-5 was obtained from Professor Joyce Loper (Oregon State University). We thank Dr Liam Elbourne (Macquarie University) for setting up our local installation of the Bio-Tradis bioinformatics pipeline, Dr Amy Cain (Macquarie University) for advice on TraDIS experimental design, Dr Francesca Short (Monash University) for advice on TraDIS data analysis, Dr Qing Yan (Oregon State University) for technical advice on the construction of knockout mutants and Professor Joyce Loper (Oregon State University) and Dr Virginia Stockwell (USDA) for valuable discussions and advice. This work was supported by an Australian Research Council Discovery Grant (DP160103746). BKF is the recipient of an Australian Government Research Training Pathway Scholarship and KAH is supported by an Australian Research Council Future Fellowship (FT180100123).

## Notes

### Competing Interest Statement

The authors have declared no competing interest.

