## Supplementary Material for "A genome-wide approach uncovers the suite of genes important for swarming motility in the biocontrol bacteria *Pseudomonas protegens* Pf-5"

**This PDF file includes:**

Table S1
Table S2
Table S3
Figure S1
Figure S2
Figure S3
Figure S4
Figure S5
References

**Other supplementary materials for this manuscript include:**

Dataset S1

Dataset S1. **Fold changes of all *Pseudomonas protegens* Pf-5 genes under swarming conditions from comparing the output (motile) pool to the control pool**. Metadata located in separate tab. Genes are considered detrimental for swarming motility fitness when their function is lost ($log_2$ fold change < -2, q < 0.01) or beneficial for swarming motility fitness when their function is lost ($log_2$ fold change > 2, q < 0.01). NA = genes excluded from the analysis as they had less than ten reads in both replicates of either the control or treatment condition.

Table S1. ***Pseudomonas protegens* Pf-5 transposon mutant library metrics from Bio-Tradis pipeline analysis**.

| Treatment | Replicate | Total no. of reads | No. (%) of reads with matching transposon tag | No. (%) of aligned reads | No. of unique insertion sites | Average distance between unique insertion sites (bp) |
|---|---|---|---|---|---|---|
| Swarming | 1 | 1,413,282 | 1,409,143 (99.7%) | 1,380,402 (98.0%) | 331,391 | 21.3 |
| Swarming | 2 | 1,478,369 | 1,473,965 (99.7%) | 1,442,536 (97.9%) | 336,380 | 21.0 |
| Control | 1 | 3,951,744 | 3,922,215 (99.3%) | 2,187,032 (55.8%) | 352,775 | 20.1 |
| Control | 2 | 2,790,059 | 2,770,318 (99.3%) | 1,941,521 (70.1%) | 345,809 | 20.5 |

Table S2. **Primers for allelic exchange mutagenesis of genes.** The random six nucleotides are italicised, restriction enzyme recognition sites are indicated in bold, and the complementary regions between the UpR/DnF primers are underlined.

| Genes | Amplification regions | Primer names | Primer sequences (5'-3') | Amino acids remaining (including start & stop codons) |
|---|---|---|---|---|
| PFL_5495 | 5' flanking | 5495-UpF-XbaI | *GTGAGG***TCTAGA**CATCCACCACTAGAATGTCC | 19 |
| | | 5495-UpR | CGACACTTTTCTTGGTTCCAGAAGACGCGTAA | |
| | 3' flanking | 5495-DnF | CTTCTGGAACCAAGAAAAGTGTCGCATCAGGT | |
| | | 5495-DnR-XbaI | *GTGAGG***TCTAGA**AAACTGAAGAATGGCCTTGC | |
| PFL_4083 | 5' flanking | 4083-UpF-XbaI | *GTGAGG***TCTAGA**TGGTCGGCGTACTCTCGGAA | 20 |
| | | 4083-UpR | GGCAATGCTCGAGTGCACTTTCATAGATCGAT | |
| | 3' flanking | 4083-DnF | ATGAAAGTGCACTCGAGCATTGCCCTGTACAAG | |
| | | 4083-DnR-XbaI | *GTGAGG***TCTAGA**TGTCCAGTGAGTAGAGCATG | |
| PFL_4091 | 5' flanking | 4091-UpF-XbaI | *GTGAGG***TCTAGA**GGAGTCGATACCGTTGTCCA | 6 |
| | | 4091-UpR | TCACGCCGGTTGGGTCATAAGTACTTCCCAGA | |
| | 3' flanking | 4091-DnF | GTACTTATGACCCAACCGGCGTGATTGTTTCC | |
| | | 4091-DnR-XbaI | *ATGACG***TCTAGA**CTTGCGCACCATGTTGATCG | |

Table S3. **Primers for verification of gene deletions via PCR and sequencing.** These primers anneal external to the gene deletion region of the genome, or within the vector backbone external to the cloned insert region.

| Genes | Functions | Primer names | Primer sequences (5'-3') |
|---|---|---|---|
| PFL_5495 | Verification of ΔPFL_5495 mutant | 5495ExtF | CAAGTCCACTACCGTTGACC |
| | | 5495ExtR | AGCCAGATGAATGCGTCGTT |
| PFL_4083 | Verification of ΔPFL_4083 mutant | 4083ExtF | GGCAACGACACCAACTTCC |
| | | 4083ExtR | TCTCGATCCTGAACCCGTAG |
| PFL_4091 | Verification of Δ*pvdE* mutant | 4091ExtF | CTGGCAGTATCAGGCGACGT |
| | | 4091ExtR | AGGTGGTGTCTTCGTTCAGG |
| pEx18Tc vector | Verification of mutant allele constructs | pEX18Tc_F | CCTCTTCGCTATTACGCCAG |
| | | pEX18Tc_R | GTTGTGTGGAATTGTGAGCG |

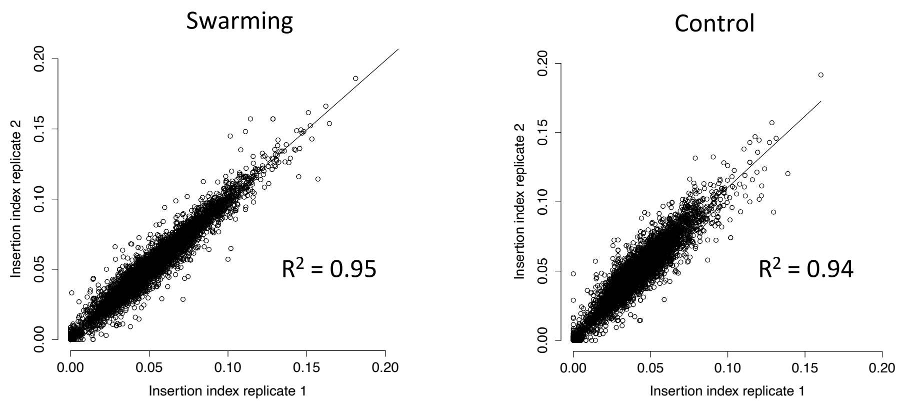


Figure S1. **Correlation of gene insertion indexes for two replicates of the *Pseudomonas protegens* Pf-5 transposon insertion library under swarming and control conditions**. Insertion index is calculated as the number of transposon insertion sites in a gene divided by the gene length. Figure generated using R(R Core Team 2018).

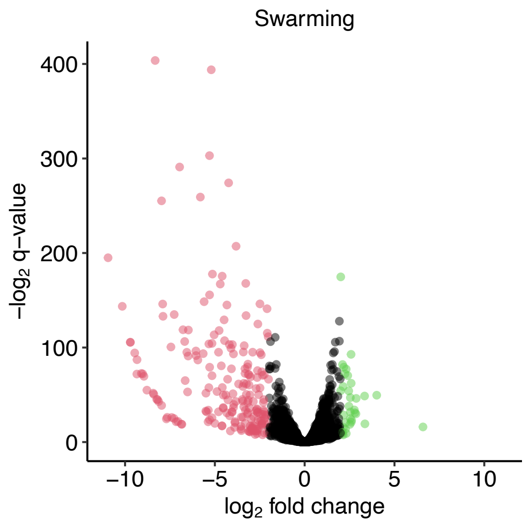


Figure S2. **Volcano plot showing $log_2$ fold change of all *Pseudomonas protegens* Pf-5 genes under swarming conditions when compared with the control.** Points in red have a significant fold change < -2 ($p < 0.01$).; loss of these genes is detrimental for swarming fitness. Points in green have a significant fold change > 2 ($p < 0.01$); loss of these genes is beneficial for swarming fitness. Figure generated using the R package ggplot2(Wickham 2016).

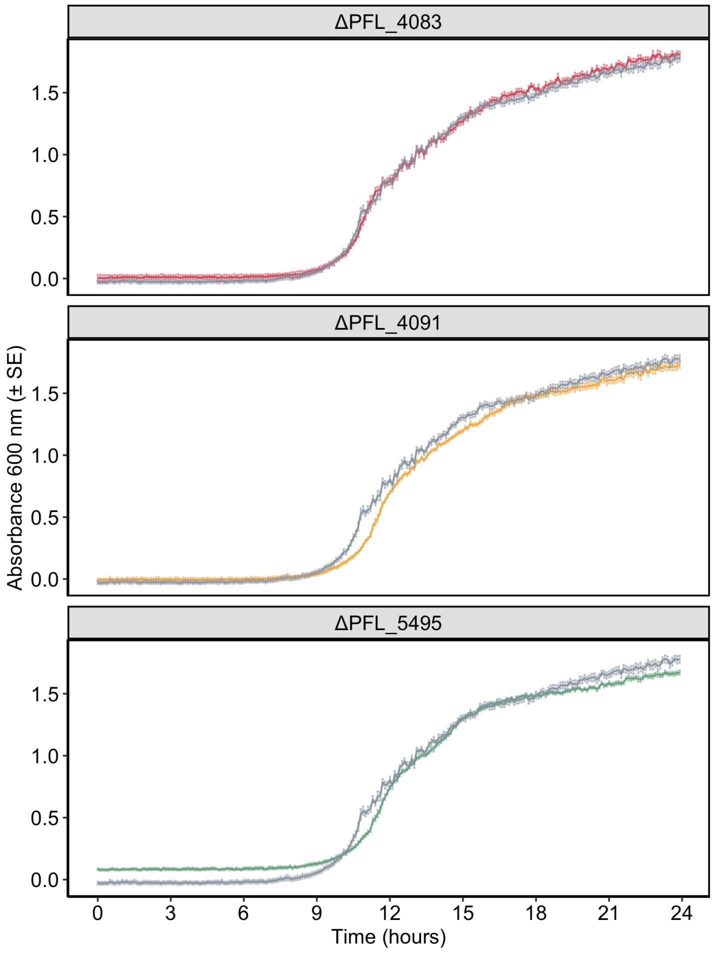


Figure S3. **Growth of *Pseudomonas protegens* Pf-5 single knockout mutants compared to wild-type (grey).** Assay conducted usi9ng cation adjusted Mueller Hinton media. Absorbance was measured at 600 nm, five replicates per strain were used and the mean absorbance (± SE) is shown. Figure generated using the R package ggplot2(Wickham 2016).

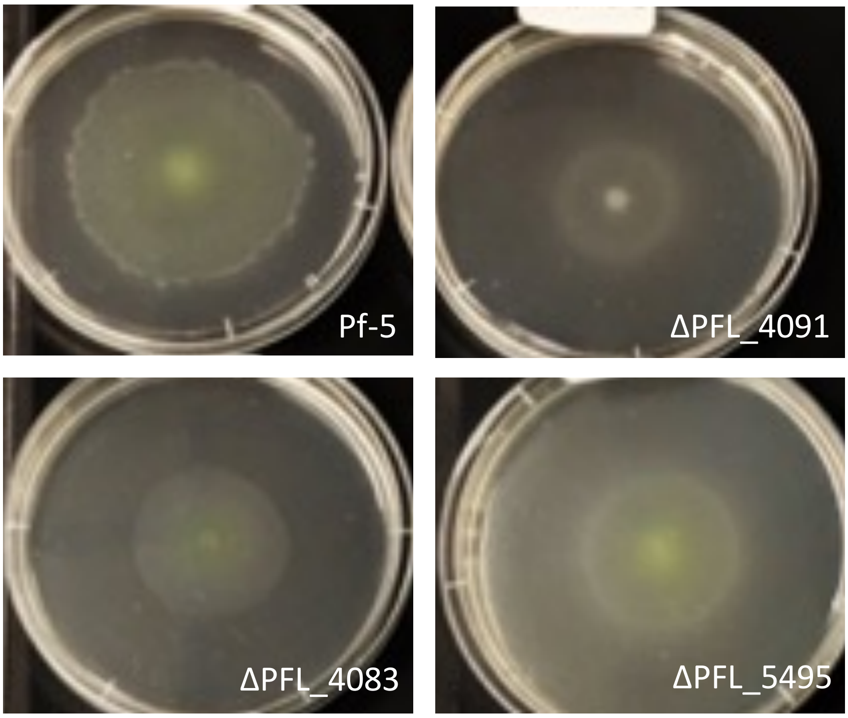


Figure S4. **Representative swarming assay plates for *Pseudomonas protegens* Pf-5 parental strain and *pvdE* (PFL_4091), PFL_4083 and PFL_5495 single knockout mutants after 24 hours growth on soft agar plates with 0.6% agar.**

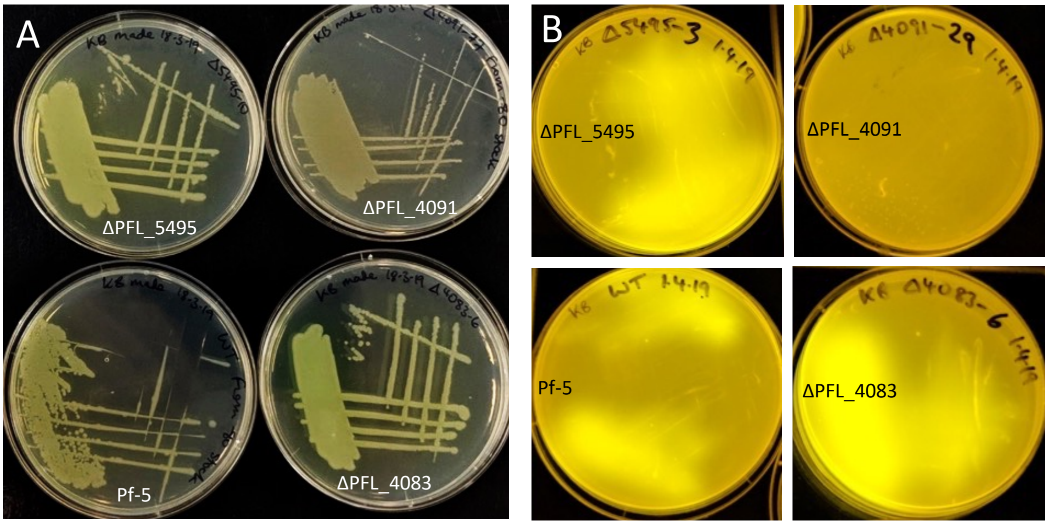


Figure S5. **Representative streak plates of *Pseudomonas protegens* Pf-5 wild-type and *pvdE* (PFL_4091), PFL_4083 and PFL_5495 knockout mutants under (A) fluorescent (white) lights and (B) UV light.** Presence of pyoverdine is indicated by yellow colouration of bacterial cells under white light and bright yellow fluorescence under UV light.
